# Macrophage migration inhibitory factor is a potential therapeutic target for cisplatin induced peripheral neuropathy in breast cancer

**DOI:** 10.1101/2025.05.13.653890

**Authors:** Hya El-Baroudy, Ximena Mejia Delgadillo, Nishi Bamania, Bhadrapriya Sivakumar, Shubham Dwivedi, Bari Chowdhury, Nickson Joseph, Snigdha Pathak, Sandra Mizkus, Shahid Ahmed, Anand Krishnan

## Abstract

**Background:** Cisplatin (CP) is an effective chemotherapy drug for several cancers. However, the use of CP is associated with peripheral neuropathy, a painful nerve disorder. Unfortunately, no therapies are available for CP-induced peripheral neuropathy (CisIPN). This study explored the role of a cytokine, the macrophage migration inhibitory factor (MIF), as a potential therapeutic target for CisIPN.

**Methods:** The role of neuroinflammation and MIF in CisIPN was evaluated in mice models of CisIPN, with and without breast cancer, after treatment with the anti-inflammatory drug Dexamethasone (Dex). Circulating MIF levels in animals were examined using ELISA. Pharmacological inhibition of MIF was achieved using the small molecule inhibitors, CPSI-1306 and ISO-1. Mechanical and thermal sensitivities of animals were assessed using von frey filament and cold acetone assays. Macrophage infiltration in peripheral nerve tissues was examined using CD68 and Iba-1 staining.

**Results:** Our results showed that Dex suppressed mechanical hyperalgesia in CisIPN animals, which was accompanied by downregulation of MIF. We also found that circulating MIF levels were increased in CisIPN animals. Furthermore, direct inhibition of MIF using CPSI-1306 and ISO-1 led to suppression of mechanical hyperalgesia, without compromising the anti-tumor efficacy of CP, in CisIPN animals. We did not find any significant change in macrophage infiltration in the peripheral nerve tissues of CisIPN animals. Immunostaining results indicated that sensory neurons in the DRGs and Schwann Cells in the sciatic nerves are potential sources for increased MIF in CisIPN.

**Interpretation:** Overall, our results strongly suggest that MIF is a promising therapeutic target for CisIPN.

## Introduction

Current advances in targeted therapies have transformed the entire landscape of cancer treatment. However, cytotoxic drugs remain as a major component of cancer chemotherapy regimens due to their high efficacy in eliminating rapidly proliferating cells. Unfortunately, their use is often associated with severe side effects. Among these, chemotherapy-induced peripheral neuropathy (CIPN) is a particularly painful complication associated with cytotoxic drugs^1, 2^. CIPN predominantly affects sensory nerves leading to abnormal and unpleasant sensations, including hyperalgesia and allodynia^2^. The incidence and severity of CIPN vary depending upon the specific chemotherapy agent, with platinum compounds, taxanes, vinca alkaloids, and bortezomib being the most well-known inducers of CIPN^3^. The unpleasant pain and nerve dysfunctions associated with CIPN often lead to premature discontinuation of the causative drug, resulting in suboptimal treatment outcomes for patients.

Cisplatin (CP) has the highest incidence of inducing CIPN among the various platinum compounds used in cancer therapy ^4^. At the same time, CP remains highly effective in various solid cancers, including the most aggressive BRCA1 mutant triple-negative breast cancer (TNBC) ^5–7^. The underlying mechanism of CP-induced peripheral neuropathy (CisIPN) is largely unknown, posing a significant challenge to the development of effective management strategies. In this study, we found that the potent anti-inflammatory drug dexamethasone (Dex) suppresses mechanical hyperalgesia associated with CisIPN. Our results further suggest that Dex-mediated suppression of hyperalgesia potentially involves the downregulation of macrophage migration inhibitory factor (MIF). MIF is a well-known cytokine implicated in pain and neuropathies associated with Guillain-Barre syndrome (GBS), diabetic neuropathy, and spinal cord injury ^8–10^. Our experiments showed that extracellular secretion of MIF is accentuated in CisIPN animal models, suggesting a direct role for MIF in CisIPN. Furthermore, using three different animal models, including a breast cancer model, and two MIF inhibitors, we demonstrated that direct inhibition of MIF suppresses mechanical hyperalgesia associated with CisIPN. Overall, our results indicate that MIF contributes to the pathogenesis of CisIPN, and therefore, it is a promising therapeutic target for CisIPN. While the exact cellular source for MIF in CisIPN is still unclear, our results suggest that excess MIF may be released from sensory neurons in the dorsal root ganglia (DRG) and Schwann Cells (SCs) in the nerves following CP treatment.

## Materials and Methods

### Breast cancer animal model

Adult female athymic nude mice (Crl:NU(NCr)-Foxn1nu) purchased from Charles River laboratories were used for breast cancer induction. Briefly, 2×10^5^ MDA-MB-231 cells suspended in 1:1 DMEM/F12 medium: Cultrex matrix (Biotechne, Cat No. 3433-010-01) were injected into either the 4^th^ or 2^nd^ mammary fat pad of animals for separate experiments. After 4 weeks of cell injections, the animals were subjected to CisIPN experiments, as described below.

### Experimental model of CisIPN

Two to five months old CDI mouse, athymic nude mouse (Crl:NU(NCr)-Foxn1nu), and an inhouse bred Cx3cr^CreERT2^-Rosa^26R-EYFP^ mouse models were used for the experiments. The parent mice for the Cx3cr^CreERT2^-Rosa^26R-EYFP^ crossbred strain, the B6.129P2(C)-*Cx3cr1^tm^*^2^.^1^*^(cre/ERT^*^2^*^)^ ^Jung^*/J and B6.129X1-Gt(ROSA)26Sor^tm1(EYFP)Cos^/J mice, were purchased from Jackson laboratories. CD1 mice were purchased from Charles River laboratories. All animal experiments were approved by the Animal Research Ethics Board (AREB) at the University of Saskatchewan. For the induction of CisIPN, the mice were given intraperitoneal injections of 2.3 mg/kg CP (Sigma-Aldrich, Cat No. 232120) for days 1-5. This was followed by 5 days of treatment interval (days 6-10) and a second cycle of CP dosing on days 11-15. The induction of neuropathy was confirmed by performing mechanical sensitivity assay.

### Measurement of plasma levels of MIF

Blood samples were collected from the submandibular vein of experimental animals. Prior to the blood collection, the animals were anesthetized using isoflurane. The blood samples were then centrifuged at 3,000 rpm for 10 minutes at 4°C to collect plasma. The plasma concentrations of MIF were measured using a commercially available MIF ELISA kit (Antibodies online, Cat No. ABIN6574190). A standard curve was generated using the recombinant MIF available in the kit and the absorbance of both standard and test samples was read at 450 nm using spectrophotometer. MIF concentrations in the test samples were calculated from the standard curve.

### Pharmacological treatment

For Dex treatment, the animals were administered 2mg/kg of Dex (Millipore Sigma, Cat No. D1756) intraperitoneally 30 minutes before CP dosing. For MIF inhibition *in vivo*, the MIF inhibitors CPSI-1306 (Cayman Chemical, Cat No. 29905) and ISO-1 (Selleckchem, Cat No. S7732) were used. The animals were intraperitoneally injected with either CPSI-1306 (20 mg/kg ip) or ISO-1 (10 mg/kg ip) 15 minutes prior to CP dosing. The control animals received saline.

### Mechanical sensitivity measurements

Mechanical sensitivity of animals was assessed using Von Frey Filament (VFF) assay. Mice were individually habituated in the testing apparatus by placing them in cubicles for 5 minutes before conducting the tests. Briefly, VFFs of varying size/number were applied to the left hind paw of each animal and the threshold stimulus required for the paw withdrawal response was recorded. A response was considered ‘positive response’ if paw withdrawal was observed three or more times out of the five trials at a specific filament size. If less than three positive responses occurred, the next filament size was tested. A minimum three baseline measurements were taken before starting CP dosing. The average of the baseline measurements was calculated for each animal to determine the mean baseline mechanical sensitivity threshold. The mechanical sensitivity recordings were continued at frequent intervals after starting CP dosing. The percentage change in mechanical sensitivity was calculated for each animal relative to their baseline sensitivity threshold.

### Cold sensitivity measurements

Cold sensitivity was evaluated in experimental animals after spraying cold acetone on their hind paw. The animals’ response to cold stimulus was then recorded as: 0 - no response, 1 - paw withdrawal, 2 - paw withdrawal + paw licking, 3 - paw withdrawal + paw licking + scratching, and 4 - jumping or peeing. Three trials were conducted for each mouse, and the average of the three trials was taken as the final reading. Multiple baseline measurements were also done for each animal before CP dosing and the average was calculated to determine the mean baseline value for each animal. The cold sensitivity measurements were continued at frequent intervals after starting CP dosing. Finally, the percentage change in cold sensitivity was calculated for each animal relative to their baseline sensitivity threshold.

### Immunohistochemistry

DRGs and sciatic nerves were isolated from the experimental animals at the end of the experiment. The isolated tissues were fixed in Zamboni’s buffer [2% paraformaldehyde (Fisher Scientific, Cat No. F79-500) and 0.5% picric acid (Ricca Chemical, Cat No.5860-16) in PBS] for 24 h at 4°C. The tissues were then washed in PBS for three times before incubating them in 20% sucrose (Fisher Scientific, Cat No. BP220-1) for another 24 h at 4°C. The tissues were mounted in optimal cutting temperature compound (OCT) and stored at ^-^80°C for a minimum 24 h before making sections. 10 µm thick sections of DRG and sciatic nerve samples were made using a cryostat and the sections were collected onto slides. The slides containing the sections were stored at −80°C for a minimum 24 h before immunostaining. For immunostaining, slides were washed in PBS for five minutes and incubated in immunoblocker [5% donkey serum (Fisher Scientific, Cat No. 56-646-05ML) and 0.3% triton-X (Millipore Sigma, Cat No. T8787) in PBS] for 30 minutes. Slides were then washed in PBS for 5 minutes and incubated with primary antibodies for 1 h at room temperature (RT). The primary antibodies used were Iba-1 (FUJIFILM Wako, Cat No. 019-19741), CD68 (Abcam, Cat No. ab53444; Abcam, Cat No. ab31630), NF200 (Millipore Sigma, Cat No. N0142), and MIF (Abcam, Cat No. ab187064). After 1h incubation, slides were washed three times in PBS for 5 minutes and then incubated with secondary antibody for another 1 h at RT. The secondary antibodies used were anti-rabbit-Alexa fluor 488 (ThermoFisher Scientific, Cat No. A-11034,), anti-mouse-Alexa fluor 488 (ThermoFisher Scientific, Cat No. A-11001), anti-mouse-Alexa fluor 647 (ThermoFisher Scientific, Cat No. A-21235), anti-rabbit-Alexa fluor 647 (ThermoFisher Scientific, Cat No. A-21245) and anti-rat-Alexa fluor 647 (ThermoFisher Scientific, Cat No. A-21247). After 1 h incubation, slides were washed in PBS for 5 minutes three times and then mounted using Diamond Antifade Mountant with DAPI (ThermoFisher Scientific, Cat No. S36973). The images were taken using Axio Observer (Zeiss) fluorescence microscope.

### Western blot

Proteins from sciatic nerves were isolated using RIPA buffer (Fisher Scientific, Cat No. PI8990) containing protease and phosphatase inhibitor cocktail (MilliporeSigma, Cat No. PI78441). Equal amount of protein was loaded for gel electrophoresis. The resolved proteins were transferred onto a PVDF membrane (Bio-Rad, Cat No. 1620177). The blocking of the membrane was done using 5% skim milk powder containing 0.1% Tween-20 (Fisher Scientific, Cat No. BP337-500) in TBS for 1 h. Following blocking, the membrane was washed twice and incubated with MIF primary antibody (Abcam, Cat No. 187064) for overnight at 4^°^C. After washing, the membrane was incubated with goat anti-rabbit HRP (Bio-Rad, Cat No.1706515) for 1h at RT. The blots were developed using an ECL substrate (Bio-Rad, Cat No. 1705060) and imaged using a GelDoc (Bio-Rad). β-actin (Fisher Scientific, Cat No. AM4302) was used as the loading control and goat anti-mouse HRP (Bio-Rad, Cat No.1706516) was used as the secondary antibody.

### Quantification of macrophage density

Macrophages were quantified in DRG and sciatic nerve sections. Macrophages in randomly selected regions (multiple regions) of the sciatic nerve or DRG sections or entire DRGs were counted using ImageJ software. The number of macrophages was then normalized to the area considered for the quantification for the calculation of macrophage density.

### Statistical Analysis

Statistical analyses were performed using the GraphPad Prism software. All data are presented as mean ± SE. Statistical tests performed included standard ‘t’ test, One-way ANOVA and two-way ANOVA with Tukey’s post hoc analysis, as appropriate. The specific details of the statistical tests performed is provided in the corresponding figure legend. p values of < 0.05 were considered statistically significant.

## Results

### Dexamethasone protects animals from CisIPN

To examine whether targeting neuroinflammation can suppress CisIPN, we treated adult male CD1 mice with the potent anti-inflammatory drug Dex before CP treatment. Animals received Dex treatment 30 minutes before CP injection for 5 days. Mechanical sensitivity of the animals was recorded using Von Frey Filaments (VFF) at frequent intervals for 8 days (**Figure S1A**). Basal sensitivity of the animals to mechanical stimuli was recorded two times before starting the CP dosing. We found a significant induction of mechanical sensitivity (mechanical hyperalgesia) in CP treated animals compared to control (saline) animals, confirming the induction of CP-induced neuropathy (**Figure S1 A, B**). Interestingly, Dex treatment suppressed CP-induced hypersensitivity remarkably, suggesting that targeting neuroinflammation can suppress CisIPN.

### Dexamethasone protects breast cancer animals from CisIPN

Evaluation of experimental CIPN therapies in cancer models is critical as the underlying cancer can influence the severity of CIPN^11^. Similarly, systemic inflammation induced by an underlying cancer can influence neuroinflammation, potentially compounding the neuroinflammation induced by chemotherapy agents^12, 13^. Thus, we evaluated the effect of Dex on CisIPN in a breast cancer animal model. Breast cancer was induced in adult female athymic nude mice by injecting MDA-MB-231 cells (TNBC cells) into the 4^th^ mammary fat pad.

At first, we examined whether breast cancer has an independent role in inducing peripheral neuropathy by recording the mechanical sensitivity thresholds of animals before and after cancer induction (**Figure 1A**). We found that animals respond to lower-size filaments after breast cancer induction, indicating that breast cancer indeed independently influences neuropathy development (**Figure 1A, B**). Next, we evaluated whether Dex could suppress CisIPN in breast cancer animals. VFF assay showed that pre-treatment with Dex could effectively suppress CP-induced mechanical sensitivity in breast cancer animals (**Figure 1C, D**). We also found a significant suppression of CP-induced cold sensitivity in breast cancer animals after pre-treatment with Dex (**Figure 1E, F**). Overall, our results show that the potent anti-inflammatory drug Dex could suppress CP-induced hypersensitivities in breast cancer animals. Moreover, there was no worsening of tumor growth in the CP+Dex combination group indicating that Dex did not compromise the tumor suppressing effect of CP (**Figure 1G, Figure S1C**).

**Figure 1:**
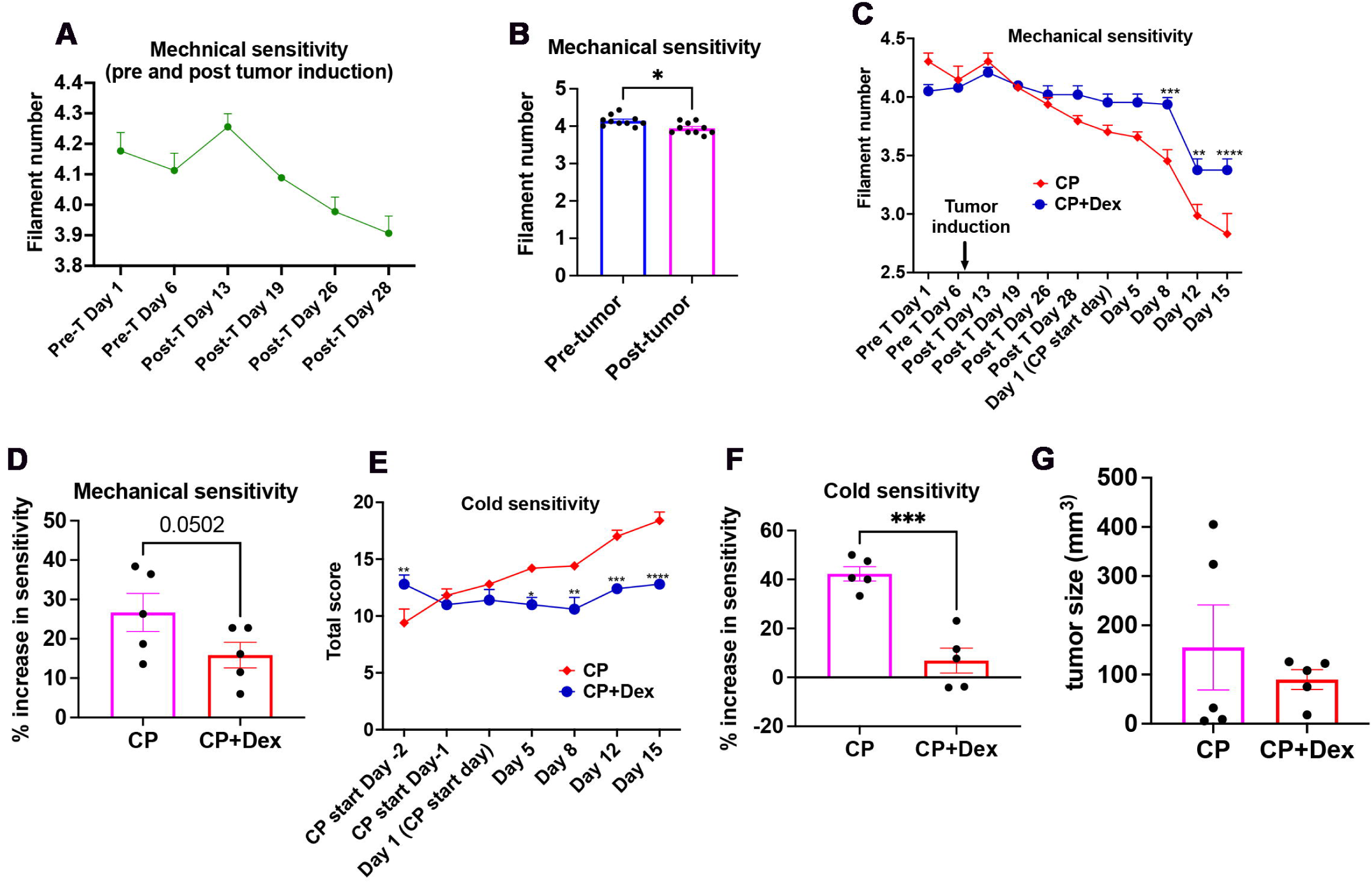
Dex suppresses CisIPN in breast cancer animals. **(A)** Line graph generated from VFF assay shows mechanical sensitivity of animals before and after breast cancer induction (Pre-T: Pre-Tumor, Post-T: Post-Tumor (n=10). **(B)** Bar graph generated from VFF assay shows mechanical hypersensitivity of animals before and after (average of days 26 and 28) breast cancer induction (mean ± SEM; n=10; paired t-test, *p<0.05). **(C)** Line graph generated from VFF assay shows suppression of CP-induced mechanical hypersensitivity in the Dex pre-treatment group [mean ± SEM; n=5; Two-way ANOVA, Sidak’s multiple comparisons test; **p<0.01, ***p<0.001, ****p<0.0001]. **(D)** Bar graph generated from VFF assay shows suppression of CP-induced mechanical hypersensitivity on Day 15 in the Dex pre-treatment group [mean ± SEM; n=5, Standard ‘t’ test; p value is shown]. **(E)** Line graph shows the sensitivity of animals to cold acetone stimuli [mean ± SEM; n=5; Two-way ANOVA, Sidak’s multiple comparisons test; *p<0.05, **p<0.01, ***p<0.001, ****p<0.0001]. **(F)** Bar graph shows the percentage increase in cold sensitivity of animals on day 15 in the indicated groups [mean ± SEM; n=5; Standard ‘t’ test; ***p<0.001)]. (**G**) Tumor volumes in CP and CP+Dex groups [mean ± SEM; n=5; Standard ‘t’ test].

### Circulating MIF levels are increased in CP treated animals

Next, to explore whether Dex-mediated suppression of CisIPN involves neuroinflammatory molecules, we surveyed the literature for neuroinflammatory molecules that are potentially linked to neuropathies and Dex. We found that the cytokine MIF contributes to pain and neuropathies associated with diabetic neuropathy, GBS, and spinal cord injuries^8–10^. Interestingly, Dex and MIF were shown to have an inverse relationship, with studies demonstrating that Dex reduces plasma levels of MIF in animal models of experimental sepsis^14^. Similarly, MIF has been shown to antagonize the growth suppressive effect of Dex in glioma cells^15^. In our study, although not statistically significant, we observed a clear trend for the downregulation of MIF in the sciatic nerves of Dex pre-treated breast cancer animals compared to CP alone treated group (**Figure S1D, E**).

No previous studies have explored the role of MIF in CIPN despite its known role in other types of neuropathies. We found that plasma levels of MIF were increased in adult female Cx3cr1^CreERT2^-Rosa^26R-EYFP^ mice after CP treatment, suggesting a potential role for MIF in CisIPN. We also noted that, compared to regular animals, the basal level of MIF in breast cancer animals was remarkably higher. For example, the basal levels of MIF in normal animals were in the range of 3 to 5000 pg/ml, while MIF levels in breast cancer animals were in the range of 5400 - 14500 pg/ml **(Figure 2 A, B (Day 0))**. We found that untreated breast cancer animals steadily declined their plasma MIF levels over time (**Figure 2B**; day 0 to day 15). However, such a decline in MIF levels was not observed in CP treated breast cancer animals, demonstrating that CP induces or maintains the levels of MIF in breast cancer animals **(Figure 2C)**. Given the potential of MIF to induce systemic inflammation and neuropathy, the high basal levels of MIF in breast cancer animals indeed suggest that MIF may contribute to cancer-induced systemic inflammation and potentially cancer-induced neuropathy.

**Figure 2:**
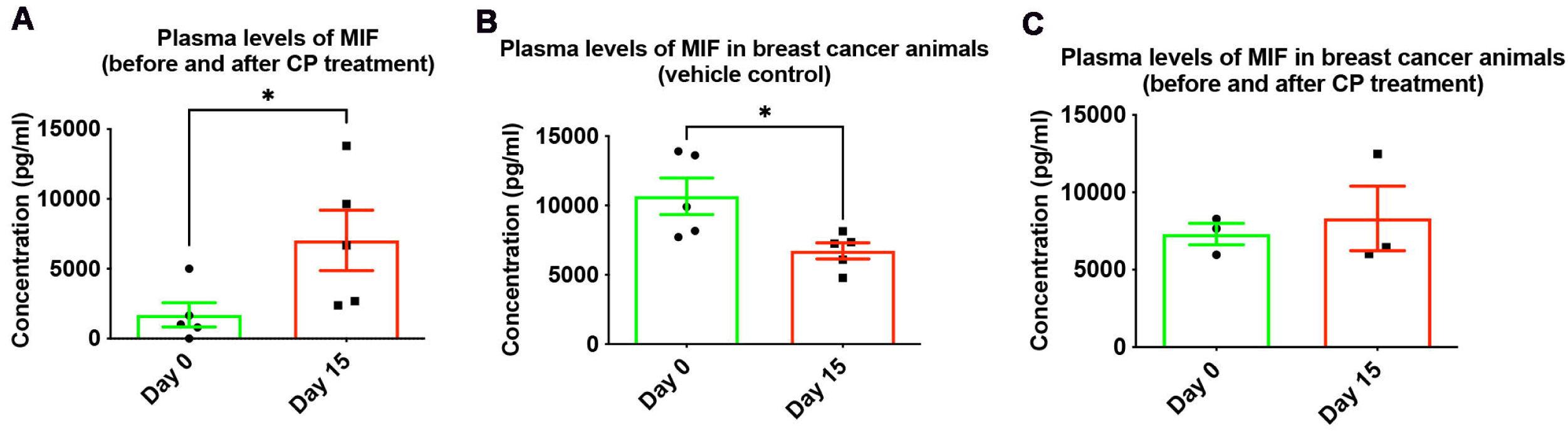
CP treatment increases circulating MIF levels. (**A**) ELISA assay shows significant induction of plasma levels of MIF after CP treatment in female Cx3cr1^CreERT2^-Rosa^26R-EYFP^ mice [mean ± SEM; n = 5; paired t-test; *p < 0.05). **(B**) ELISA assay shows plasma levels of MIF in breast cancer animals before (Day 0) and after (Day 15) vehicle treatment [mean ± SEM; n = 5; paired t-test; *p < 0.05]. (**C**) ELISA assay shows plasma levels of MIF in breast cancer animals before (Day 0) and after (Day 15) CP treatment [mean ± SEM; n = 3; paired t-test].

### Inhibition of MIF protects animals from CP-induced mechanical hypersensitivity

Our results showed that MIF is elevated after CP treatment, suggesting that MIF may contribute to CisIPN. Therefore, we examined whether inhibition of MIF could protect animals from CisIPN. Mechanical sensitivity experiments in Cx3cr1^CreERT2^-Rosa^26R-EYFP^ mice showed that the MIF the inhibitor CPSI-1306 significantly suppresses CP-induced mechanical hypersensitivity, indicating that MIF inhibition protects animals from CisIPN (**Figure 3A,B**). To check the reproducibility of this result, we repeated the experiments in CDI mice induced with CisIPN. Two MIF inhibitors, CPSI-1306 and ISO-1, were used to validate the potential of MIF as a therapeutic target for CisIPN. Consistent with previous experiments, the CP-treated group showed a decrease in the mechanical threshold, while the control, individual MIF inhibitors (CPSI-1306 or ISO-1), and the combination groups (CP+CPS1-1306 and CP+ISO-1) maintained relatively stable mechanical thresholds throughout the study period, substantiating that MIF inhibitors could protect animals from CP-induced mechanical hypersensitivity (**Figure 3 C, D**).

**Figure 3:**
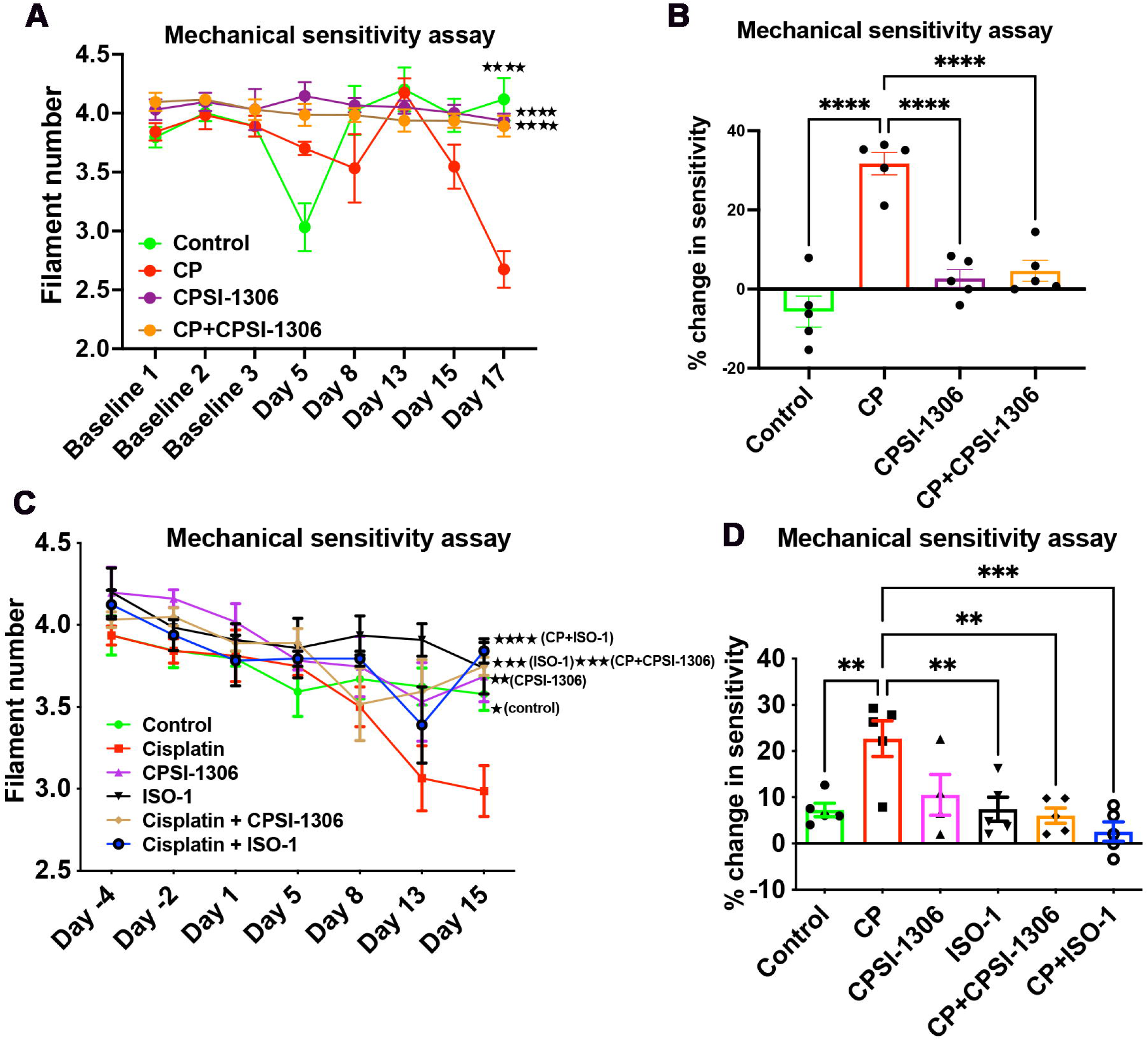
Inhibition of MIF suppresses mechanical hypersensitivity in experimental models of CisIPN. **(A)** Line graph generated from VFF assay shows mechanical sensitivity of animals (female Cx3cr1^CreERT2^-Rosa^26R-EYFP^ mice) in the indicated groups [mean ± SEM; n=5; Two-way ANOVA, Tukey’s multiple comparisons test; ****p<0.0001 (significance shown is compared to CP group)]. **(B)** Bar graph generated from VFF assay shows that pre-treatment with the MIF inhibitor CPSI-1306 significantly suppresses CP-induced mechanical hypersensitivity in CisIPN animals (female Cx3cr1^CreERT2^-Rosa^26R-EYFP^ mice) on Day-17 [mean ± SEM; n= 5; One-way ANOVA with Tukey’s multiple comparisons test; ****p < 0.0001). **(C)** Line graph generated from VFF assay shows mechanical sensitivity of animals (female CD-1 mice) in the indicated groups [mean ± SEM; n=5 (n=4 for CPS1-1306 group); Two-way ANOVA, Tukey’s multiple comparisons test; *p<0.05, **p<0.01, ***p<0.001, ****p<0.0001 (significance shown is compared to CP group)]. **(D)** Bar graph generated from VFF assay shows that pre-treatment with MIF inhibitors, CPSI-1306 and ISO-1, significantly suppresses CP-induced mechanical hypersensitivity in CisIPN animals (CD-1 mice) on Day-15 [mean ± SEM; n= 5 (n=4 for CPS1-1306 group); One-way ANOVA with Tukey’s multiple comparisons test; **p<0.01, ***p<0.001].

### MIF inhibition has no influence on CP-induced mild cold allodynia

We next examined whether CisIPN also involves cold allodynia and if MIF inhibition influences the same. The paws of animals were stimulated using cold acetone to evaluate the cold sensitivity of animals. The behavioural responses of animals to the cold stimulus were ranked on a scale from 0 to 4 (0: - no response, 1: paw withdrawal, 2: paw withdrawal and paw licking, 3: paw withdrawal, paw licking, and scratching, and 4: jumping or peeing). The experiments were performed in adult female CD-1 mice. Although there was a mild trend for increased cold sensitivity in CP treated animals, it was not statistically significant, indicating that CP does not significantly induce cold allodynia (**Figure S2 A, B**). While pre-treatment with ISO-1 appears to suppress the mild cold allodynia induced by CP, it was not statistically significant (**Figure S2B)**.

### Inhibition of MIF protects breast cancer animals from CP-induced mechanical hypersensitivity

We next evaluated the effect of MIF inhibition on CisIPN in breast cancer animals. Breast cancer was induced in adult female athymic nude mice by injecting MDA-MB-231 cells into the second mammary fat pad. As expected, the mice responded to lower-size filaments after breast cancer induction, substantiating that breast cancer has an independent role in inducing neuropathy (**Figure 4A, B**). Further, to evaluate the effect of MIF inhibition on CisIPN in breast cancer animals, mechanical sensitivity measurements were done using VFF assay. As expected, CP induced mechanical hyperalgesia in breast cancer animals as evident from the animals’ response to lower mechanical thresholds compared to saline treated control breast cancer animals. Interestingly, pre-treatment of animals with MIF inhibitors (CPSI-1306 or ISO-1) suppressed the CP-induced mechanical hyperalgesia, however, the effect was significant only for CPSI-1306, suggesting that CPSI-1306 may be more effective in relieving CisIPN in breast cancer animals (**Figure 4 C, D**).

**Figure 4:**
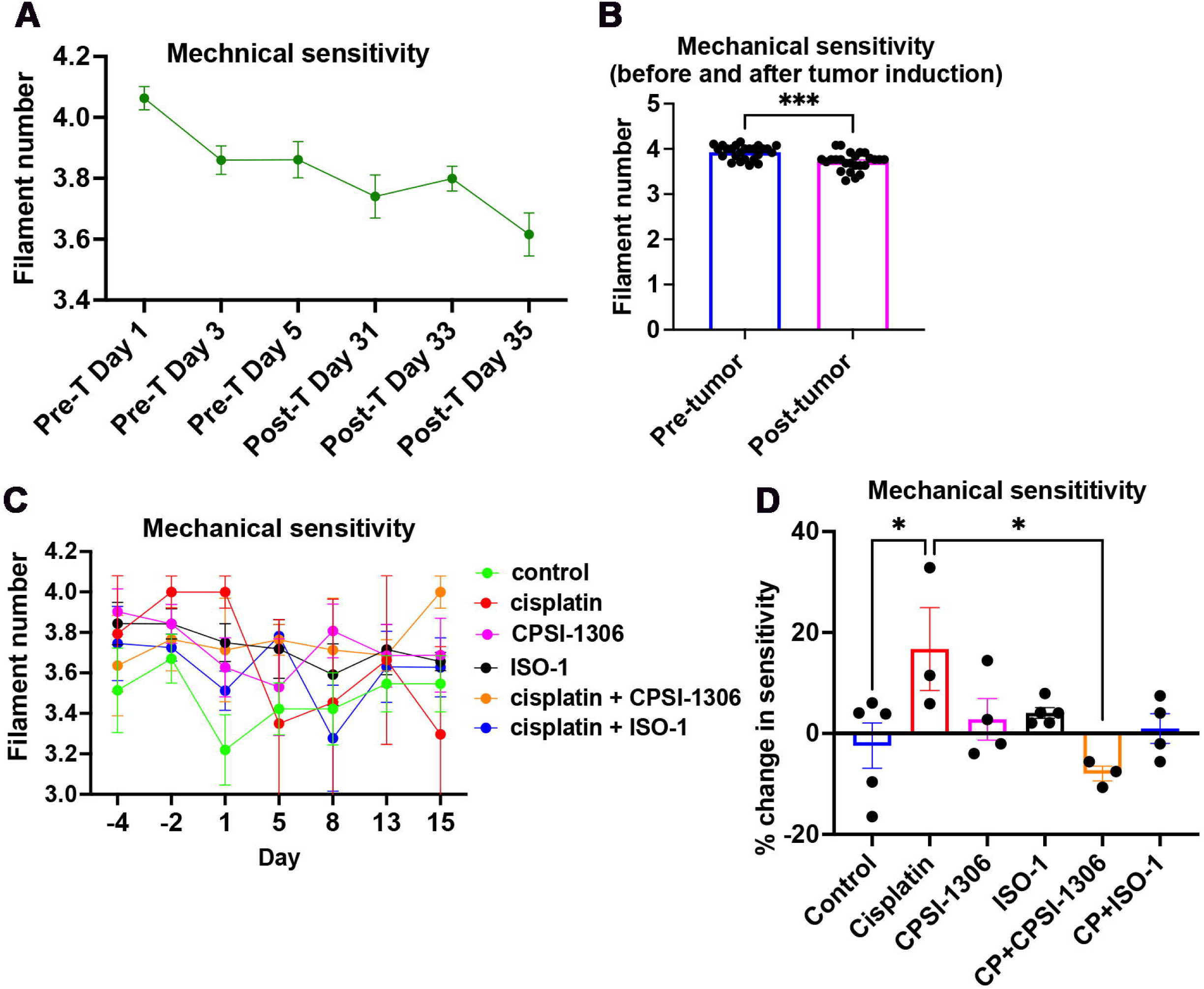
**(A)** Line graph generated from VFF assay shows mechanical sensitivity of animals before and after breast cancer induction (Pre-T: Pre-Tumor, Post-T: Post-Tumor) [mean ± SEM; n=25]. **(B)** Bar graph generated from VFF assay shows mechanical hypersensitivity of animals before and after (day 35) breast cancer induction (mean ± SEM; n=25; paired t-test, ***p<0.001). **(C)** Line graph generated from VFF assay shows mechanical sensitivity of breast cancer animals in the indicated groups [mean ± SEM; n=5 (control and ISO-1 groups), n=4 (CPSI and CP+ISO-1 groups), n=3 (CP and CP+CPSI groups); Two-way ANOVA, Tukey’s multiple comparisons test]. **(D)** Bar graph generated from VFF assay shows that pre-treatment with CPSI-1306 significantly suppresses CP-induced mechanical hypersensitivity in breast cancer animals on Day-15 [mean ± SEM; n=5 (control and ISO-1 groups), n=4 (CPSI and CP+ISO-1 groups), n=3 (CP and CP+CPSI groups); One-way ANOVA with Tukey’s multiple comparisons test; *p<0.05].

We observed high variabilities in tumor responses in this experiment. Notably, some animals treated with CP, ISO-1, or the combination of CP+ISO-1 exhibited complete tumor regression after 15 days of treatments. Importantly, MIF inhibitors did not worsen tumor progression. Rather, the MIF inhibitors, especially when used in combination with CP, suppressed tumor progression, indicating the potential therapeutic use of MIF inhibitors for CisIPN in breast cancers (**Figure S3**).

### CP does not significantly influence macrophage infiltration in the DRGs and sciatic nerve of experimental animals

Infiltration of immune cells, including macrophages, into nerve tissues is a salient feature of neuroinflammation. Therefore, we examined the infiltration of macrophages in the DRGs and sciatic nerves of CisIPN models generated from adult female CDI mice to investigate whether CisIPN involves neuroinflammation and if MIF inhibitors suppress this process. We used the two well-established macrophage markers Iba-1 and CD68, as reported previously, to label macrophages^16, 17^. The total number of macrophages was normalised to the area of the tissue analyzed. A minimum of three biological replicas were considered per experimental group. The final data is expressed as the relative fold change in macrophage density compared to the control. Interestingly, we found no significant differences in the abundance of Iba1^+^ and CD68^+^ macrophages in DRGs and sciatic nerves of animals in the experimental groups compared to the control, indicating that CisIPN may not involve significant macrophage recruitment to peripheral nerve tissues, and thus, may not significantly involve neuroinflammation **(Figure 5 & Figure 6).**

**Figure 5:**
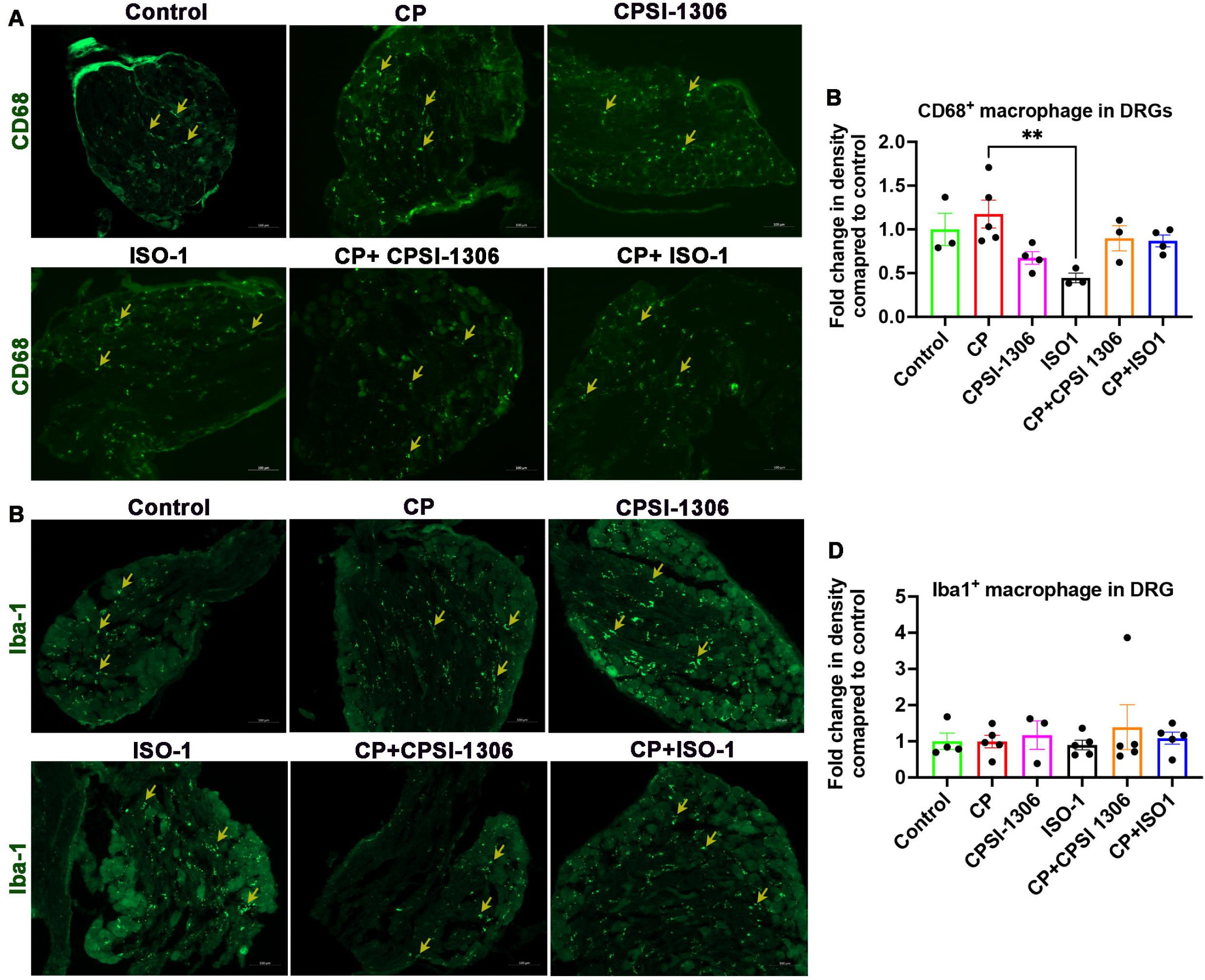
**(A)** Representative images of CD68^+^ macrophages (yellow arrows) in the DRGs of experimental animals (female CD1 mice) in the indicated groups (scale bar, 100µm). (**B**) Bar graph shows relative fold change in CD68^+^ macrophage density in the DRGs of experimental animals (female CD1 mice) in the indicated groups (mean ± SEM; n=5 (CP), n=4 (CPSI-1306, CP+ISO-1), n=3 (control, ISO-1, CP+CPSI-1306); One-way ANOVA with Tukey’s multiple comparisons test). **(C)** Representative images of Iba-1^+^ macrophages (yellow arrows) in the DRGs of experimental animals (female CD1 mice) in the indicated groups (scale bar, 100µm). (D) Bar graph shows relative fold change in Iba-1^+^ macrophage density in the DRGs of experimental animals (CD1 mice) in the indicated groups (mean ± SEM; n=5, except control (n=4) and CPSI-1306 (n=3); One-way ANOVA with Tukey’s multiple comparisons test).

**Figure 6:**
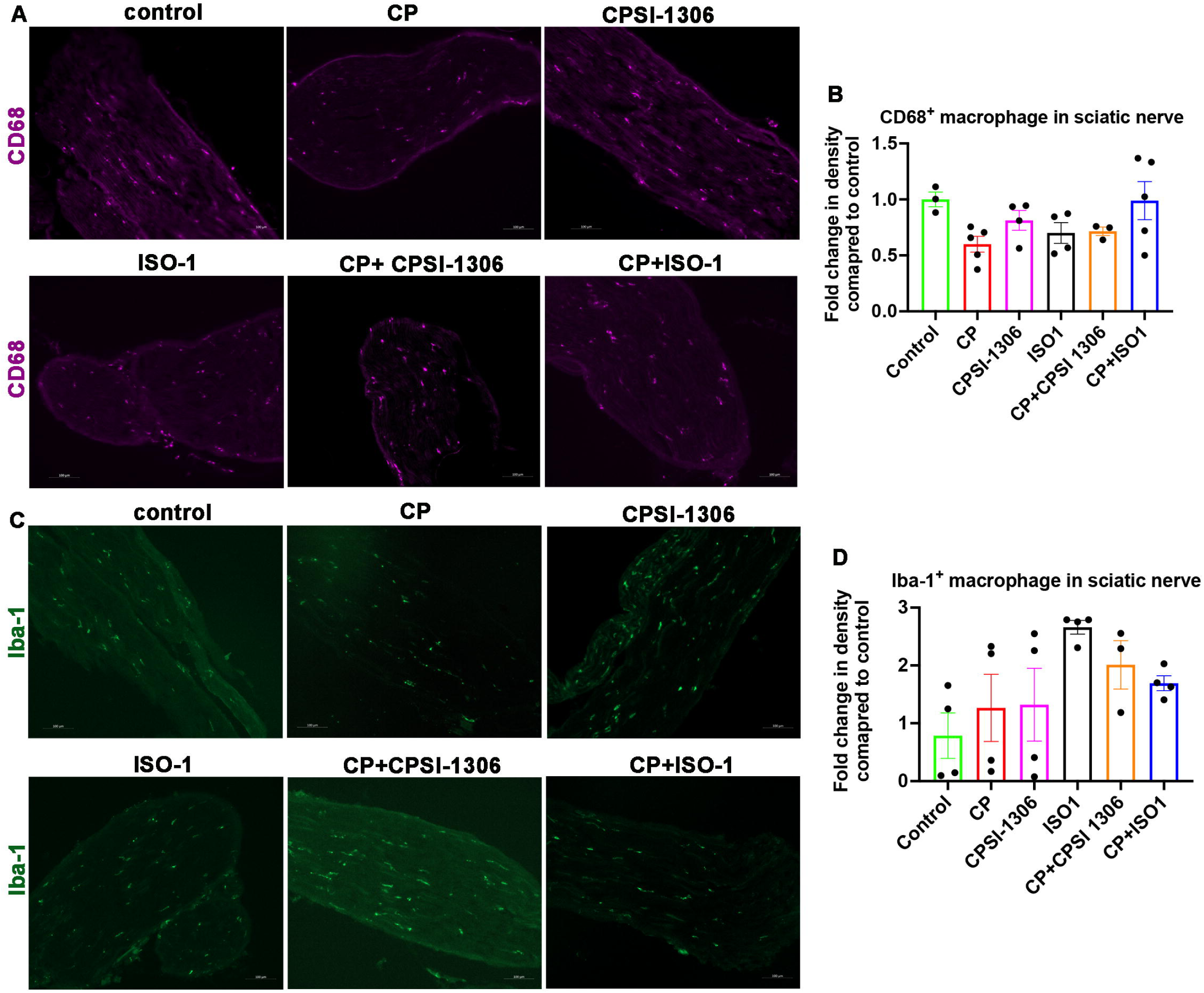
**(A)** Representative images of CD68^+^ macrophages (yellow arrows) in the sciatic nerve of experimental animals (female CD1 mice) in the indicated groups (scale bar, 100µm). (**B**) Bar graph shows relative fold change in CD68^+^ macrophage density in the sciatic nerve of experimental animals (female CD1 mice) in the indicated groups (mean ± SEM; n=5 (CP, CP+ISO-1), n=4 (CPSI-1306, ISO-1), n=3 (control, CP+CPSI-1306); One-way ANOVA with Tukey’s multiple comparisons test). **(C)** Representative images of Iba-1^+^ macrophages (yellow arrows) in the sciatic nerve of experimental animals (female CD1 mice) in the indicated groups (scale bar, 100µm). (**D**) Bar graph shows relative fold change in Iba-1^+^ macrophage density in the sciatic nerve of experimental animals (female CD1 mice) in the indicated groups (mean ± SEM; n=4, except CP+CPSI-1306 (n=3); One-way ANOVA with Tukey’s multiple comparisons test).

### Sensory neurons in the DRGs and non-axonal cells in the peripheral nerves are primary sources for MIF in CisIPN

Since we noted no significant macrophage infiltration in the DRGs and sciatic nerves of CP treated animals, we speculated additional cellular sources for MIF in peripheral nerves. Immunostaining experiments showed that MIF is preferentially expressed in NF200^poor^ small calibre neurons, which mediates pain transmission, in the DRGs of untreated breast cancer animals (**Figure 7A**). We also noted a gain of expression of MIF in NF200^high^ large calibre neurons after CP treatment in the breast cancer model, indicating that sensory neurons in the DRGs is a chief source for MIF in CisIPN (**Figure 7A**). In the sciatic nerve, although MIF was weakly expressed by axons, it was most predominantly expressed by SCs, as evident from our CD1 mice studies (**Figure 7B, C**). We also observed an intense expression of MIF in nerve resident SCs after CP treatment, suggesting that SCs are the primary source for MIF in sciatic nerves following CisIPN induction.

**Figure 7:**
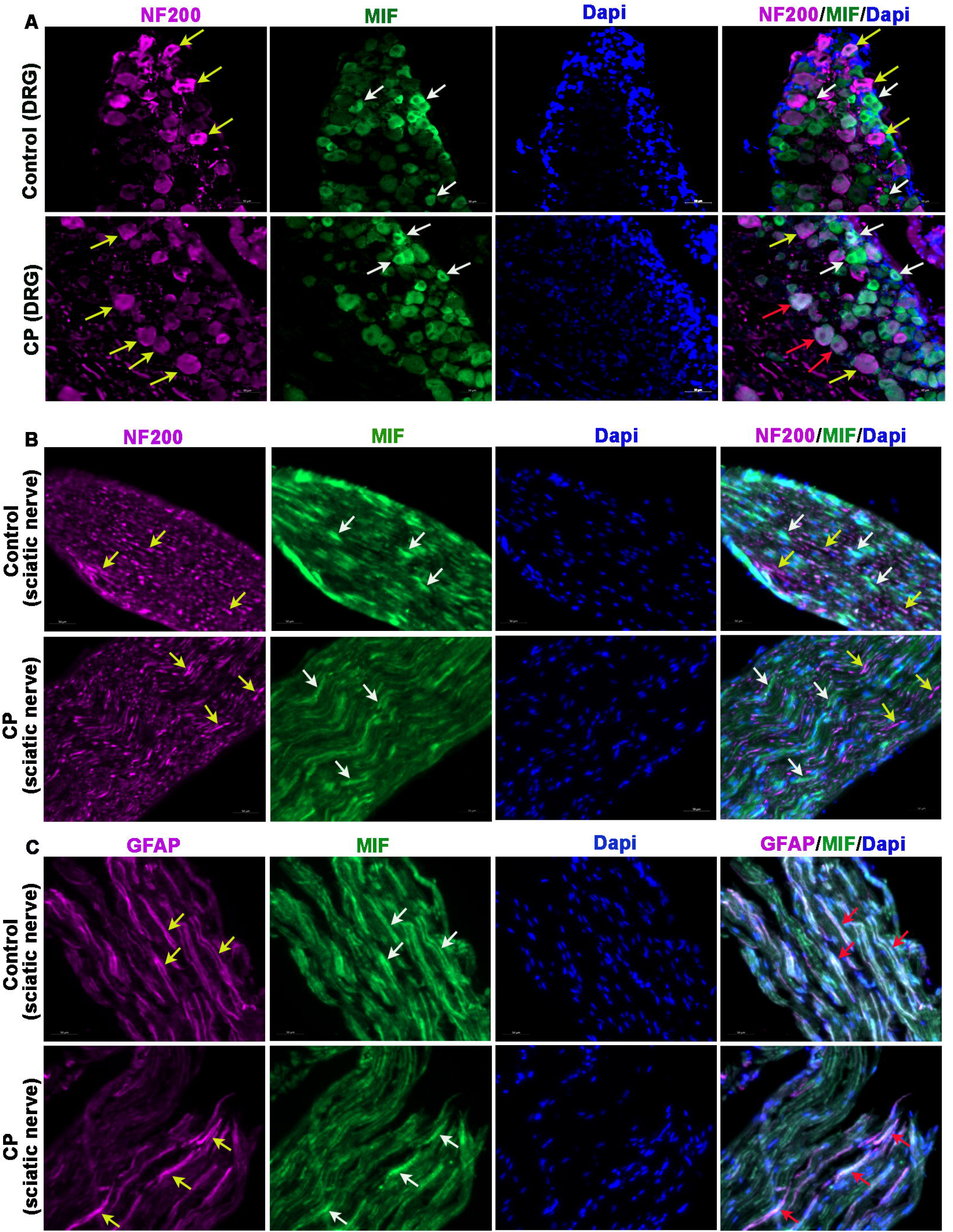
**(A)** Representative images of NF200 (yellow arrows) and MIF (white arrows) in the DRGs of control (saline) and CP treated animals (breast cancer model). Co-localization of NF200 and MIF is shown by red arrows in the merged CP group panel (scale bar, 50µm). (**B**) Representative images of NF200 (yellow arrows) and MIF (white arrows) in the sciatic nerves of control (saline) and CP treated animals (female CD1 mice). (**C**) Representative images of GFAP (yellow arrows) and MIF (white arrows) in the sciatic nerves of control (saline) and CP treated animals (female CD1 mice). Co-localization of GFAP and MIF is shown by red arrows in the merged panel (scale bar, 50µm).

## Discussion

Despite being a debilitating side effect of cancer chemotherapy, no effective treatment is currently available to completely reverse CIPN other than discontinuing or reducing the dose of the causative chemotherapy agent^18^. Several causative mechanisms have been suggested for CIPN in the past, however, none of them have proven to be effective intervention points for developing CIPN therapies. Moreover, the presence of unique CIPN mechanisms associated with individual classes of chemotherapeutic agents further complicate the identification of effective therapies. This work is specifically focused on CisIPN. Previous reports suggest that neuroinflammation can potentially influence CisIPN progression^13^. Surprisingly, we did not find any change in macrophage infiltration in nerve tissues, a hallmark of neuroinflammation, after CP treatment, even though we observed that the potent anti-inflammatory drug Dex could suppress CisIPN. This indicates that CisIPN may involve neuroinflammation driven by resident macrophages or neuroinflammatory molecules released by other cell types, such as neurons and glial cells. Thus, a mild suppression of MIF we observed in peripheral nerves after Dex treatment in our CisIPN models supports the critical role of MIF derived from either neurons, resident macrophages, and/or glial cells in inducing and/or facilitating CisIPN.

We found that circulating MIF levels are elevated in mice after CP treatment. Individuals with spinal cord injuries were shown to have increased circulating levels of MIF^10^. Patients with diabetic neuropathy also showed increased MIF levels in their footpad lesions^9^. Additionally, MIF levels were shown to elevate in the cerebrospinal fluids of multiple sclerosis patients^19^. All these pathological conditions are associated with neuropathy, substantiating the involvement of MIF in inducing neuropathies. Here, for the first time, we demonstrated the involvement of MIF in CisIPN.

We considered regular (*Cx3cr1^CreERT^*^2^*-Rosa^26R-EYFP^ and CD1 mice)* and tumor (TNBC) bearing mouse models for this study. The regular animal models are rapid screening models, while tumor models provide insights into the role of MIF in CisIPN under the influence of a solid cancer. The Cx3cr^CreERT2^-Rosa^26R-EYFP^ model expresses YFP in macrophages upon tamoxifen administration. However, we used tamoxifen non-injected animals and performed immunostaining to visualize macrophages for a head-to-head comparison of macrophages across the experimental models used in this study. CP is a well-established therapy for TNBC, supporting the relevance of the cancer model we used in this study^7, 20^. Importantly, cancer itself can induce neuropathy^11^. The emerging evidence of a crosstalk between solid tumors and peripheral nerves support the independent roles of cancers in inducing neuropathy^21–24^. Thus, our observation of elevated levels of MIF in tumor bearing mice compared to healthy mice strongly suggests a role for MIF in facilitating cancer-induced neuropathy. TNBC patients were shown to express higher levels of MIF in tumors compared to adjacent normal tissues, and this observation further supports our arguement^25^.

Mechanical hypersensitivity and cold allodynia are two well-known measurable indices established for CIPN. Previous studies showed that direct injection of recombinant MIF into mice produces a dose-dependent increase in mechanical sensitivity, indicating that MIF can independently promote mechanical hyperalgesia^26^. Interestingly, our results consistently demonstrated that pharmacological inhibition of MIF using CPSI-1306 and ISO-1 effectively suppresses CP-induced mechanical hyperalgesia, providing strong evidence that MIF facilitates neuropathic pain in CisIPN. Overall, these results are encouraging and suggest MIF inhibitors as potential therapies for CisIPN. Previous studies showed the potential of MIF inhibitors in suppressing neuropathic pain in other pathological conditions. For example, pharmacological inhibition of MIF was shown to suppress pain-like behaviors in animal models of nerve injury^27^. Our experiments did not show reproducible induction of cold allodynia after CP treatment across the experimental models. A previous study also showed that CP is weak in inducing cold allodynia compared to other platins, such as oxaliplatin, which aligns with the observations we made in this study^28^.

Although unexpected, we did not find significant changes in macrophage infiltration in nerves and DRGs after CP treatment, suggesting that recruited macrophages may not be the primary source for MIF in CisIPN. A recent study also showed that CP has no major influence in inducing macrophage infiltration nerves, matching the observations we made here^29^. We evaluated macrophage infiltration after 10 doses of CP, and hence, we do not know if macrophage infiltration occurs at early dosing periods. Regardless, the lack of macrophage filtration in nerves and DRGs posed questions on the source of elevated circulating MIF in our CisIPN models. We observed that MIF is expressed by sensory neurons in the DRGs with a sub-population of large calibre sensory neurons gaining MIF expression after CP treatment.

Similarly, we found that the major cellular constituents of peripheral nerves, including axons and SCs express MIF, with SCs expressing MIF most predominantly at the basal and CP treated conditions. This suggests that sensory neurons in the DRGs and nerve-resident SCs in the nerves are the primary source for elevated circulating MIF in CisIPN. However, additional investigations are warranted to assess whether macrophages, especially a sub-population of self-renewing nerve-resident macrophages, also induce circulating MIF in CisIPN^30^.

We observed high variability in the final tumor volumes of animals in the breast cancer experiments (MIF inhibitor experiments), with some animals in the CP, ISO-1, and CP+ISO-1 groups regressed tumors completely after 15 days, challenging the statistical representation of the final tumor volumes. Regardless, we did not find tumor progression in MIF inhibitor groups compared to the control, rather it appeared that MIF inhibition independently and in combination with CP suppressed the tumor growth. This finding suggests that MIF-based therapies potentially do not worsen tumor burden when used in cancer patients. Previous studies showed that MIF promotes solid cancers^25, 31^. In line with this, inhibition of MIF was also previously shown to suppress cancers^25, 32, 33^. Thus, anti-MIF approach in CisIPN may offer dual benefit by relieving neuropathy and suppressing cancer growth.

Overall, this study has identified, for the first time, that MIF contributes to CisIPN. Our findings suggest that targeting MIF represents promising therapeutic strategy, especially for alleviating the mechanical hyperalgesia component of CisIPN. These encouraging results warrant further investigation of MIF inhibitors in male models of CisIPN and across other forms of CIPN.

## Author contributions

HEB performed experiments, analyzed data, and contributed to preparing the initial version of the manuscript. XMD, NB, BS, SD, BC, NJ, SM, and SP contributed to data generation and analysis. SA contributed to data analysis and edited the manuscript. AK conceived the concept, provided the resources for the work, analyzed the data, prepared the initial and revised versions of the manuscript, and finalized the manuscript.

## Data availability

The data supporting this work is included in the article. No data belonging to this article is deposited in shared repositories.

## Ethics statement

The animal experiments described in this work have been approved by the Animal Research Ethics Board (AREB) at the University of Saskatchewan.

## Supplementary figures

**Figure S1:**
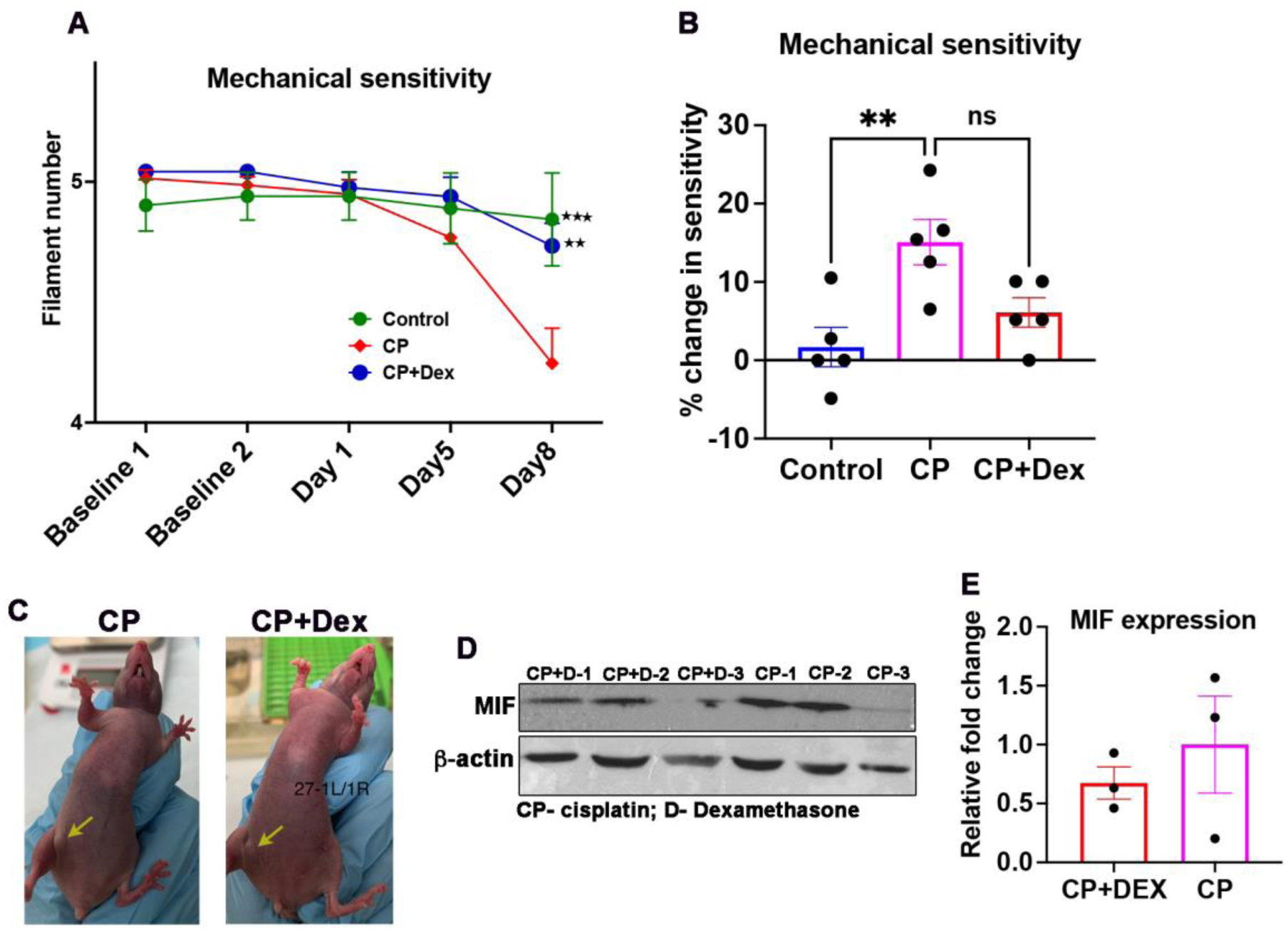
Dexamethasone suppresses cisplatin-induced peripheral neuropathy. **(A)** Line graph generated from Von Frey Filament (VFF) assay shows mechanical sensitivity of animals (male CD1 mice) in the indicated groups [mean ± SEM; n=5; Two-way ANOVA, Tukey’s multiple comparisons test; **p<0.01, ***p<0.001 compared to CP group]. **(B**) Percentage change in mechanical sensitivity of animals (male CD1 mice) on day 8 compared to the basal mechanical sensitivity [mean ± SEM; n=5; One-way ANOVA, Tukey’s multiple comparisons test; **p<0.01]. (**C**) Representative images of tumors in CP and CP+Dex treated animals. **(D)** Western blot shows the expression of MIF in the indicated experimental groups. β-actin was used as the loading control. (**E**) quantification of ‘C’ [mean ± SEM; n=3; Standard ‘t’ test].

**Figure S2:**
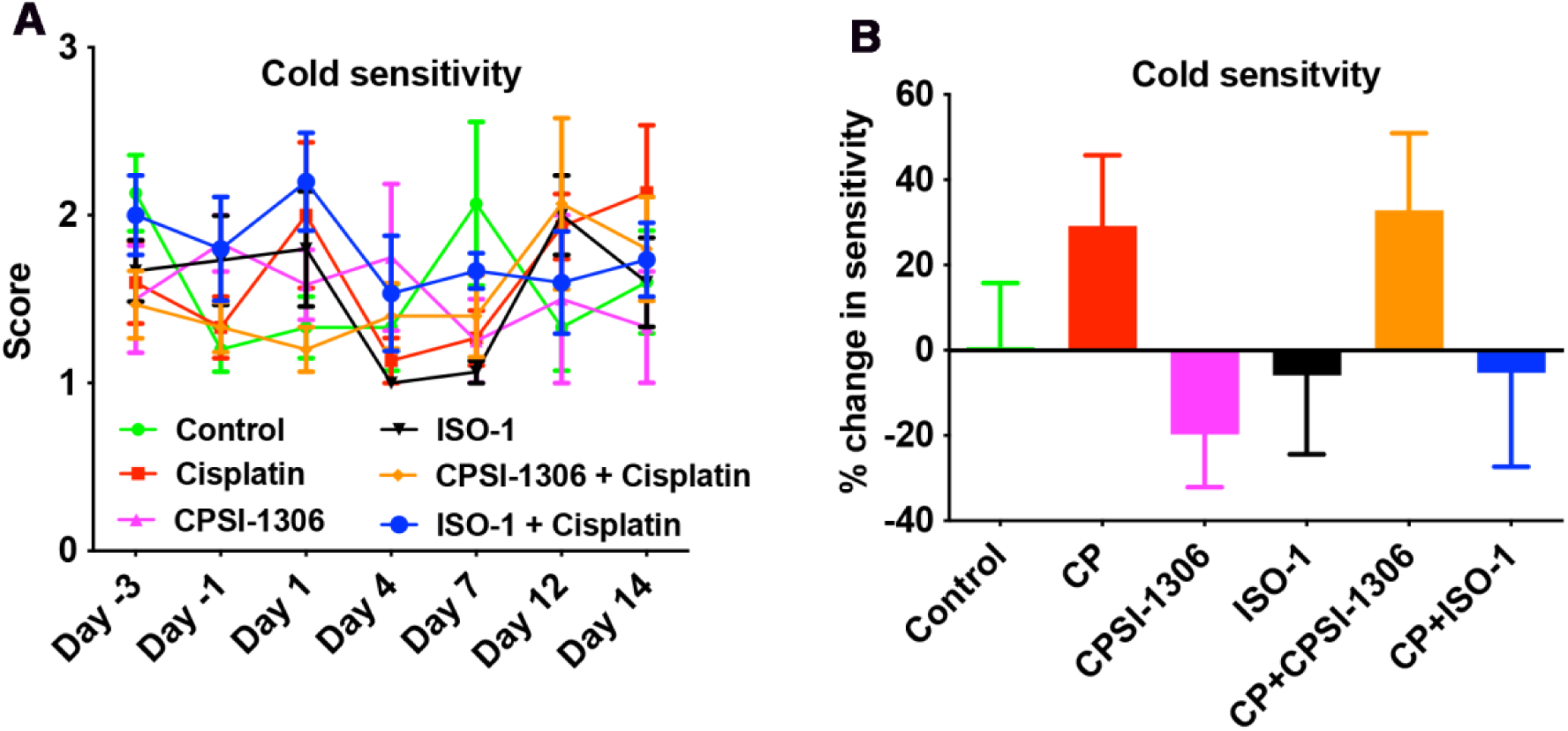
Cold sensitivity measurements in CD1 mice. (**A**) Line graph generated from cold acetone assay shows the cold sensitivity of animals in the indicated groups [mean ± SEM; n=5 (n=4 for CPS1-1306 group); Two-way ANOVA, Tukey’s multiple comparisons test]. (**B**) Bar graph generated from cold acetone assay shows mild induction of cold allodynia in CP group on Day-14, while pre-treatment with ISO-1 appears to suppress the same (not statistically significant) [mean ± SEM; n= 5 (n=4 for CPS1-1306 group); One-way ANOVA with Tukey’s multiple comparisons test].

**Figure S3:**
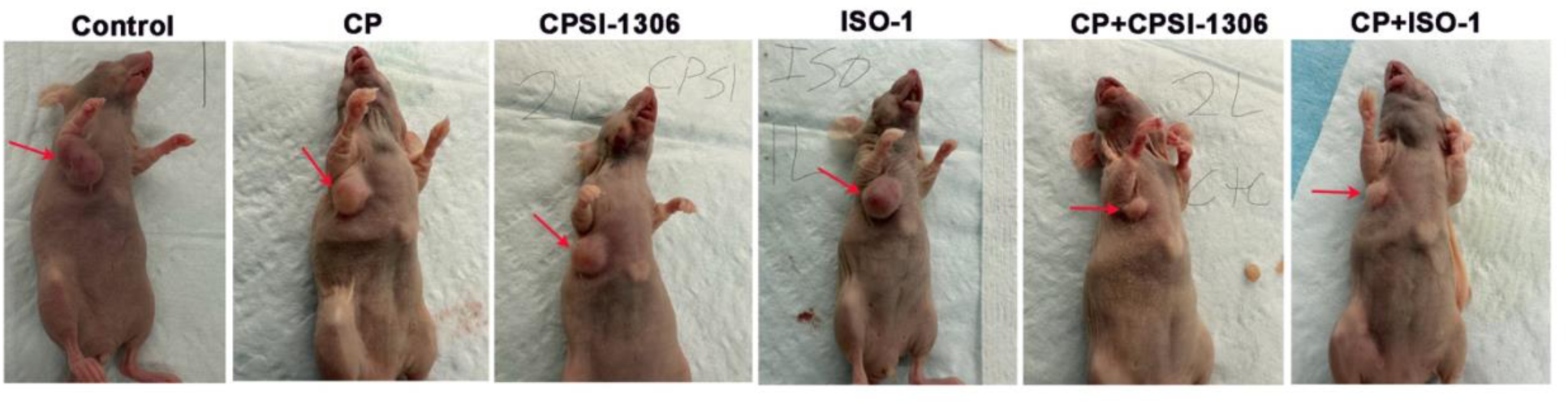
Representative images of breast tumors (red arrows) in the experimental animals.

